# IGF2BP3 inhibits IL-13 and IL-4 effects in human airway epithelium and is dysregulated in type 2 disease

**DOI:** 10.1101/2025.07.19.665669

**Authors:** Paniz Khooshemehri, Onofrio Zanin, Jennifer Rynne, Esperanza Perucha, Rocio T Martinez-Nunez

## Abstract

Type 2 immunity encompasses the coordinated cellular responses mainly driven by IL-4 and IL-13 to target extracellular parasites. Uncontrolled, type 2 responses underpin asthma and allergy. At the cellular level, IL-13 and IL-4 bind to receptors IL-13Rα1 and IL-4R and trigger STAT6 phosphorylation leading to the transcription of type 2 mediators. Further regulators of this fundamental pathway remain poorly understood. We demonstrate that the RNA binding protein Insulin Growth Factor 2 Binding Protein 3 (IGF2BP3) inhibits IL-13/IL-4 effects in airway epithelium and is increased in type 2 pathology, where it correlates with an IL-13-driven signature. Mechanistically, IGF2BP3 directly binds *IL4R* and *IL13RA1* mRNAs, which are also methylated. Depleting IGF2BP3 increased *IL4R* and *IL13RA1* mRNAs half-life, IL-4Rα and IL-13Rα1 surface expression and IL-13/IL-4-dependent STAT6 phosphorylation. Reducing IGF2BP3 levels skewed primary airway epithelial cells towards a type 2 phenotype and enhanced IL-13-mediated transcriptional effects genome-wide. Lastly, we found IGF2BP3 levels upregulated in airway epithelium in several type 2 disease cohorts, and an IGF2BP3-dependent IL-13-driven signature predominant in type 2 high vs type 2 low asthma. Our data positions IGF2BP3 as a novel inhibitor of IL-13/IL-4 effects and type 2 disease biomarker.

## Introduction

Type 2 immune responses are designed to protect the body against parasites such as tape worms. Uncontrolled, they drive the pathology of allergic diseases, asthma, dermatitis, or ulcerative colitis (1). As exemplar of type 2 disease where IL-4 and IL-13 are key disease drivers (2, 3) , patients with are classified into type 2 high vs type 2 low according to clinical parameters including fractional exhaled nitric oxide (FeNO), atopy and eosinophilia (4). Type 2 immunity comprises the broad immune response that is mainly driven by the archetypal cytokines IL-4 and IL-13, together with IL-5, IL-31 and IL-9 (1). IL-4 and IL-13 share surface receptor subunits that can be arranged into two types of receptors. Type I receptors are composed of IL-4R and common gamma chain (γc) and Type II receptors are composed of IL-4R and IL-13Rα1. While IL-4 can bind Type I and II receptors, IL-13 solely binds to Type II receptors (5). Additionally, IL-13 can also bind to IL-13Rα2 which appears to have distinct signalling properties (6). IL-4 and IL-13 share signalling pathways, mainly driven by the phosphorylation of signal transducer and activator of transcription 6 (STAT6), triggering transcriptional responses that drive the mRNA expression of multiple mediators, including eotaxins, alarmins and interleukins (reviewed in (5). However, IL-4 and IL-13 also have distinct and non-overlapping functions in a cell-dependent manner, possibly due to differential receptor expression. While Type I receptors are expressed in multiple cell types, Type II receptors are restricted to innate immune cells including airway epithelial cells (5).

The airway epithelium sits at the tissue-environment interface, serving as both barrier and regulator of tissue homeostasis. Epithelium is thus the first barrier that encounters pathogens and environmental challenges. The epithelium is disrupted, both physically and phenotypically, in multiple chronic inflammatory conditions on the rise, characterised by excessive IL-13/IL-4-driven processes (1, 7). For example, IL-4 and IL-13 trigger increased mucus production, as well as chemokine and eotaxin expression in epithelium, creating a pro-inflammatory milieu that perpetuates type 2-driven inflammation in the underlying mucosa.

IL-4 and IL-13 responses have been extensively investigated at the transcriptional level, but post-transcriptional regulation remains poorly understood. Post transcriptional regulation encompasses all processes that modulate the fate of a given RNA after transcription, from 5’ capping, 3’ tailing, splicing, chemical modification, stabilization, transport or translation (8, 9). RNA binding proteins (RBPs) can modulate each one of these steps, while microRNAs mainly modulate the stability and translation of their target mRNAs (10). We and others have previously shown that microRNAs can modulate IL-13-driven responses (11–13). RBPs are known to contribute to the inflammatory response, mainly via downregulation of inflammatory mediators including cytokine-encoding genes (8, 14). While most models focus on type 1 responses (driven by mediators such as tumour necrosis factor alpha (TNF-α), IL-1 or interferons) (15), the role of RBPs in IL-13/IL-4 responses remains unknown.

Insulin Growth Factor 2 Binding Protein 3 (IGF2BP3) is an RBP that belongs to the family of IGF2BPs together with IGF2BP1 and IGF2BP2. Initially described as only present during embryogenesis (16), IGF2BP3 is significantly upregulated in various tumours and tumour-derived cell lines (Yaniv & Yisraeli, 2002; Yisraeli, 2005) and is thus considered an oncofoetal factor (17). Upon binding to its target RNA, IGF2BP3 can regulate RNA stability, localisation, and translation, as well as mRNA-microRNA interactions (18, 19). IGF2BP3 is also a N(6)-methyladenosine reader (20), and can promote mRNA translation and stability (20) or switch target mRNAs from translating polyribosomes to non-translating intracellular bodies (21). Thus, IGF2BP3 appears to have cell type-dependent molecular functions.

Interestingly, IGF2BP3 can activate the JAK/STAT pathway (22), typically associated with cancer and cytokine signalling (23). We observed IGF2BP3 levels upregulated in airway epithelial cells from patients with asthma (24), and searched for possible targets that were immune- and airway-related in publicly available RNA immunoprecipitation (RIP) datasets. We found data suggesting that IGF2BP3 binds to transcripts encoding *IL4R* and *IL13RA1* in pancreatic cells (25). These findings led us to hypothesise that IGF2BP3 modulates *IL4R* and *IL13RA1* mRNA fate and/or expression and IL-13/IL-4 signalling consequently. These regulatory processes might participate in the maintenance of lung epithelium homeostasis (1, 26), and when dysregulated contribute to type 2 disease. We therefore tested the role of IGF2BP3 in IL-13/IL-4 signalling in human airway cells, its potential as a novel modulator of IL-13/IL-4-driven responses and investigated IGF2BP3 levels and effects in type 2 disease.

## Results

### IGF2BP3 is present in adult healthy human lung and binds IL4R and IL13RA1 transcripts

We had previous evidence that *IGF2BP3* mRNA is present in human bronchial epithelium and bound to polyribosomes – indicating possible translation (24), despite IGF2BP3 being previously described as a foetal (16) or oncogenic factor (22). We tested IGF2BP3 presence in different primary cells as controls, including peripheral blood leukocytes (CD14^-^ cells, CD14^+^ cells and granulocytes) considering their prominent role in immunity. We also tested the expression of IGF2BP3 protein in primary bronchial epithelial cell extracts exposed to dexamethasone or control where we had previously explored the presence of other RBPs (27). While we found low-to-undetectable levels of IGF2BP3 in blood leukocytes (Figure 1A, lanes 1-3), we found clear immunoreactivity of IGF2BP3 in primary bronchial epithelial cells, despite loading half the protein amount as compared to blood cells (Figure 1A, samples 1-4).

**Figure 1.**
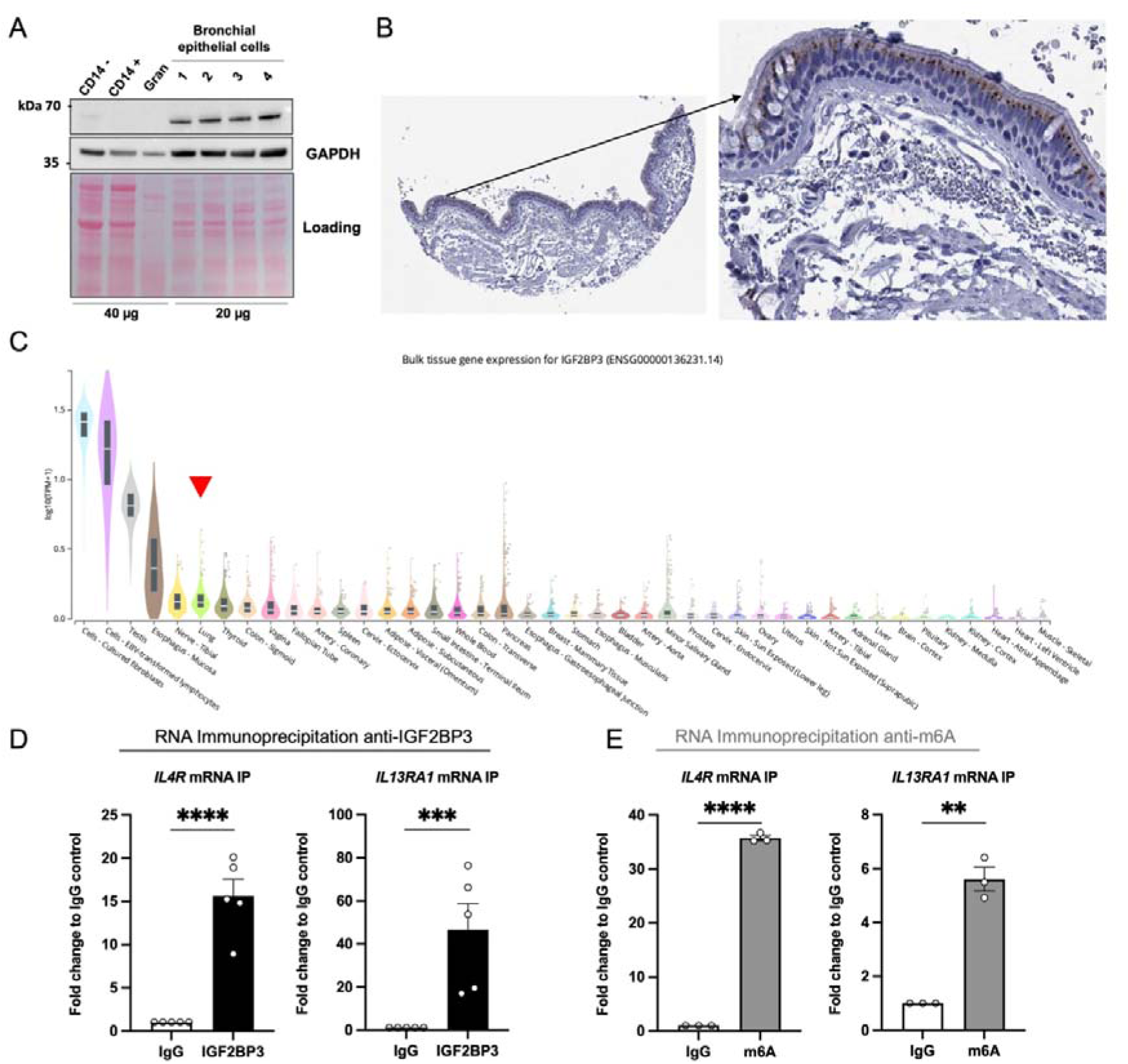
IGF2BP3 is expressed in healthy human adult lung epithelium, and it binds *IL4R* and *IL13RA1* mRNAs. A: Western blot image of lysates from leukocyte fractions (lanes 1-3) and healthy cultured human bronchial epithelial cells exposed to dexamethasone (samples 2 and 4) or vehicle (samples 1 and 3) from (27). B: Immunohistochemistry representative image showing presence of IGF2BP3 protein in lung epithelium. Image credit: Human Protein Atlas, www.proteinatlas.org, (28). C: GTEx expression plot for *IGF2BP3* mRNA showing normalised counts per human tissue. The red triangle marks lung. D: RNA immunoprecipitation of IGF2BP3 or IgG control showing enriched precipitation *IL4R* and *IL13RA1* mRNAs (n=5). E: RNA immunoprecipitation of m(6)A or IgG control showing enriched precipitation *IL4R* and *IL13RA1* mRNAs. Data (n=5; n=3) depicted as mean ± SEM. Gran: granulocytes. supp analysis was done using a two-tailed ratio test. * *P* ≤ 0.05; ** *P* ≤ 0.01; *** *P* ≤ 0.001; **** *P* ≤ 0.0001.

Considering that our primary bronchial epithelial cells had been placed in culture, we also interrogated public databases for *ex vivo* and *in situ* levels of IGF2BP3. Figure 1B depicts an immunohistochemistry image of IGF2BP3 in human lung from the Human Protein Atlas (28), where IGF2BP3 can be observed in epithelium, particularly on the apical side. Using GTEx (29) we also found *IGF2BP3* mRNA preferentially present in certain cells and tissues, with the top being cultured fibroblasts and EBV-transformed lymphocytes. Amongst tissues, lung had the 4^th^ highest expression (marked with a red triangle), after testes, oesophageal mucosa and tibial nerve (Figure 1C). We found a wide range of expression and also multiple tissues with little or absent *IGF2BP3* mRNA levels (Figure 1C; for the full set from GTEx see Supplementary Figure 1). Using GTEx single cell datasets, we also observed enrichment of IGF2BP3 expression in epithelial cells in multiple tissues, including lung (Supplementary Figure 2).

To determine IGF2BP3 functionality, we searched previous RNA immunoprecipitation data and found that IGF2BP3 binds to *IL4R* and *IL13RA1* transcripts in a pancreatic cell line (25). Considering the importance of IL-13/IL-4 signalling in bronchial epithelium (1, 26), we assessed whether IGF2BP3 can bind *IL4R* and *IL13RA1* mRNAs in bronchial epithelial cells. We found endogenous IGF2BP3 significantly immunoprecipitated *IL4R* (*P* <0.0001) and *IL13RA1* (*P* = 0.0003) mRNAs as compared to IgG control (Figure 1D) in BEAS-2B cells. Because IGF2BP3 is a N(6)-methyladenosine (m6A) reader (20, 21) we tested if *IL4R* and *IL13RA1* are methylated mRNAs. Indeed, we found *IL4R* and *IL13RA1* co-precipitated with m6A immunocomplexes as compared with IgG control (*IL4R P* <0.0001, *IL13RA1 P* = 0.002, Figure 1E). Together, our data show that IGF2BP3 is expressed in human lung and airway epithelium, where *IL4R* and *IL13RA1* are methylated RNAs that are bound by IGF2BP3.

### IGF2BP3 depletion causes an increase in mRNA half-life and protein surface expression of IL4R and IL13Rα1

To determine the effects of IGF2BP3 on *IL4R* and *IL13RA1* mRNA expression, we generated constitutively depleted IGF2BP3 cell lines using shRNA, and concomitant scrambled control cell lines (*P* = 0.0052, Figure 2A). IGF2BP3-knocked down cells did not have different levels of *IL4R* or *IL13RA1* mRNAs (*P* = 0.9976 and *P* = 0.7683, respectively, Figure 2B). However, we found increased expression of IL-4R (*P* = 0.0246) and IL-13Rα1 (*P* = 0.0116) on the surface of IGF2BP3-depleted cells (Figure 2C). Considering that IGF2BP3 can modulate the stability of its bound transcripts (30), we exposed shIGF2BP3 vs shControl BEAS-2B cells to actinomycin D and measured the mRNA levels of *IL4R* and *IL13RA1* using RT-qPCR over time. We found a mild increased mRNA half-life for *IL4R* (123.2 vs 101.1 minutes) and tripled for *IL13RA1* (65.53 vs 21.25 mins) in IGF2BP3-depleted BEAS-2B cells when compared to control. Thus, IGF2BP3 depletion increases IL-4R and IL-13Rα1 surface expression at least partially by increasing their encoding mRNA half-lives.

**Figure 2.**
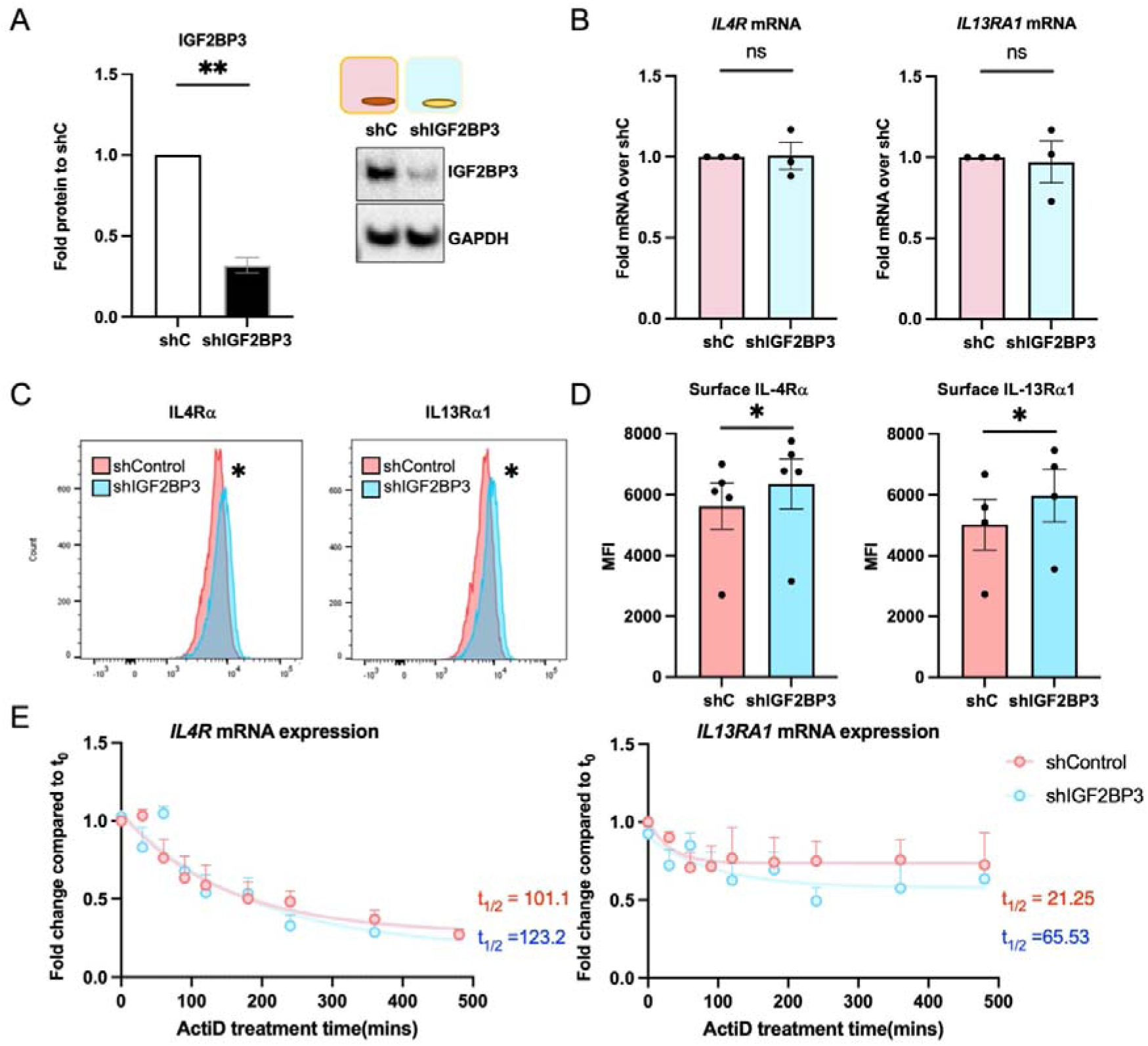
IGF2BP3 depletion leads to increased IL-4R and IL-13Rα1 surface expression and mRNA stability. A: Representative western blot and quantification of IGF2BP3 depletion in IGF2BP3 shRNA cell lines (shI) vs scrambled RNA control (shC). B. *IL4R* and *IL13RA1* mRNA levels in shRNA BEAS-2B cell lines. C. Histograms depicting IL-4R and IL-13Rα1 surface expression and (D) quantification of median fluorescence intensity (MFI) of IL-4Rα and IL-13Rα1 in shControl and shIGF2BP3 BEAS-2B cells. Statistical analyses were done using two-tailed t-tests on log transformed data. E: mRNA stability analysis of *IL4R* and *IL13RA1* mRNA expression on IGF2BP3-depleted cells vs scrambled control. Fold changes were fitted to a non-linear regression and one-phase exponential decay analysis to determine half-lives as per (31). Data (n=3-5) depicted as mean ± SEM. Two-tailed ratio t-test unless differentially stated. * *P* ≤ 0.05; ** *P* ≤ 0.01.

### IGF2BP3 depletion leads to increased IL-4/IL-13-dependent signalling

IL-4 and IL-13 trigger STAT6 phosphorylation to increase the transcription of IL-4/IL-13/STAT6-dependent genes (5). Concomitant with the increased levels of IL-4R and IL-13Rα1 (Figures 2C and 2D) in our constitutive cell lines, we found subtle but statistically significantly increased phosphorylation of IL-4- (*P* = 0.042) and IL-13-triggered (*P* = 0.0051) STAT6 phosphorylation in IGF2BP3 depleted BEAS-2B cells (Figure 3A) 1h after stimulation. This effect was enhanced downstream, as we observed increased levels of the IL-4- and IL-13-dependent mRNAs *CCL11* (IL-4 *P* = 0.0313 and IL-13 *P* = 0.03) and *GATA3* (IL-4 *P* = 0.0226 and IL-13 *P* = 0.0073) (Figure 3B), and a nearly significant increase (*P* = 0.058) of IL-13-dependent *CCL26* mRNA. Intriguingly, we also observed increased mRNA levels of the transcription factor *GATA3* (32) (*P* = 0.0101) in an IL-4- and IL-13-independent manner. At the protein level, we found increased IL-13-driven secretion of TSLP (*P* = 0.0342), CCL11 (*P* = 0.0439) and IL-8 (*P* = 0.021) (Figure 2C). Depletion of IGF2BP3 therefore leads to increased IL-4 and IL-13-dependent STAT6 phosphorylation, accompanied by an increase in mRNA expression and secretion of type 2 mediators.

**Figure 3.**
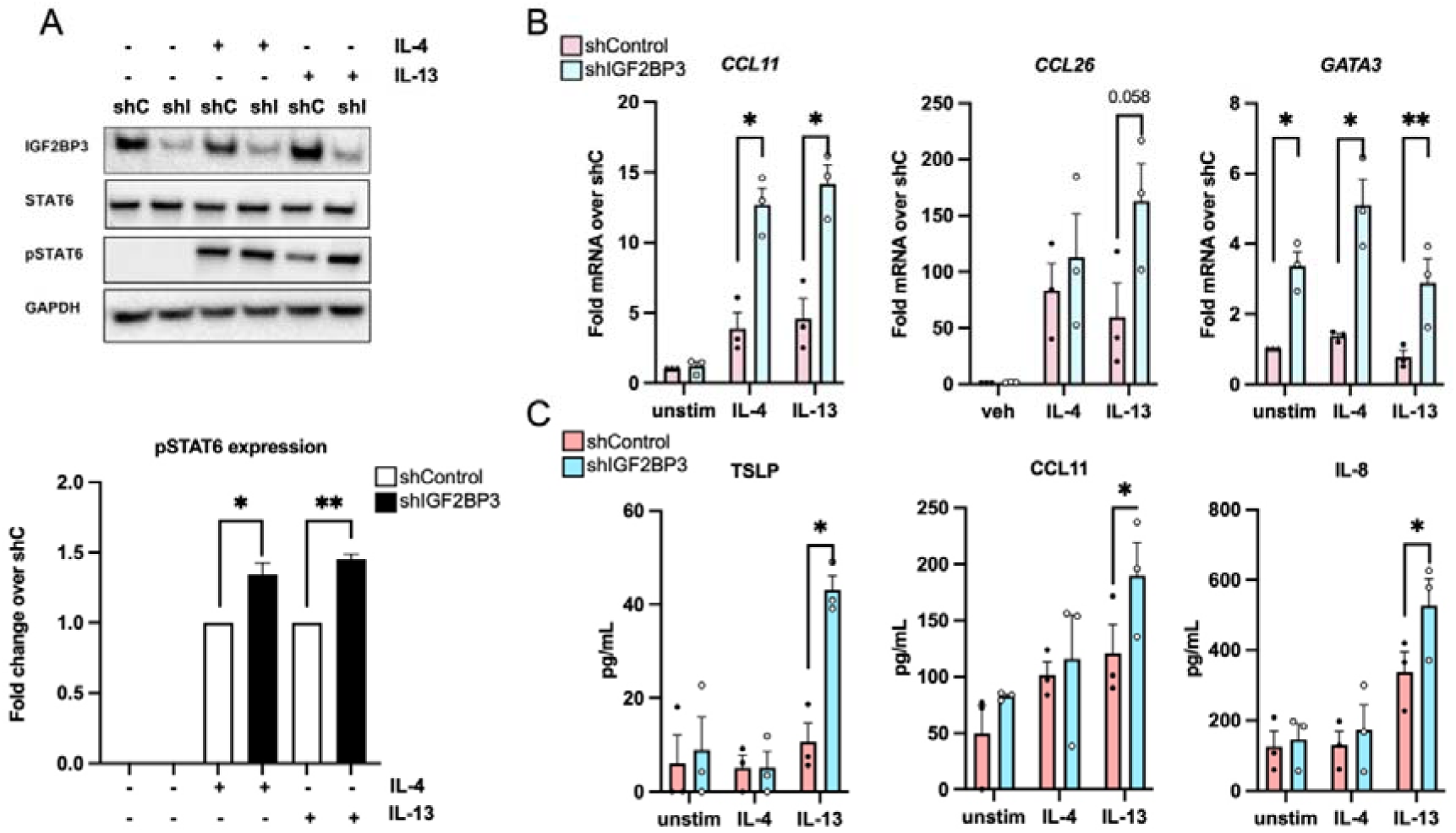
IGF2BP3 depletion leads to increased IL-4/IL-13-dependent signalling. A: top: representative western blot of BEAS-2B cells constitutively depleted of IGF2BP3 (shIGF2BP3) or scrambled RNA control (shControl), stimulated with IL-13 or IL-4 (1 hour). Bottom: Densitometry quantification. Statistics were done using two-tailed ratio tests. B: mRNA levels of type 2 mediators in shIGF2BP3 vs shControl cells, in the presence of IL-4, IL-13 or control vehicle. Statistics were done using two-tailed ratio tests. C: quantification of secreted proteins shIGF2BP3 vs shControl cells, in the presence of IL-4, IL-13 or control vehicle. Data (n=3) depicted as mean ± SEM. Statistics were done using two-tailed t tests. * *P* ≤ 0.05; ** *P* ≤ 0.01. unstim: unstimulated (referring to vehicle stimulated).

### IGF2BP3 depletion in primary bronchial epithelial cells increases their IL-4- and IL-13-dependent responses

BEAS-2B are widely utilised in the field of airways research, but they are nonetheless a cell line immortalized with SV40 virus (33). To analyse a more physiological system, we investigated whether IGF2BP3 depletion would alter IL-13/IL-4-driven mRNA transcription in primary bronchial epithelial cells from healthy donors. IGF2BP3 depletion was also successful in primary human cells (Figure 4A tope left panel, unstimulated *P* = 0.0025, IL-4 *P* = 0.0005 and IL-13 *P* = 0.0002). We observed very subtle but statistically significant changes in the main mediators of IL-13/IL-4 signalling upon IGF2BP3 knock down (Figure 4A), namely *IL13RA1* and *STAT6* mRNAs (*P* = 0.0098, *P* = 0.0217, respectively) as well as in the presence of IL-4 (*P* = 0.0048, *P* = 0.0057 respectively) but not IL-13. *GATA3* mRNA levels were also increased upon IGF2BP3 knock down in the presence of IL-4 (*P* = 0.0132). IL-4-and IL-13-driven mRNA expression of type 2 mediators was also enhanced (Figure 4B). We focused on chemokines that attract leukocytes and eosinophils, noting that primary BECs express different chemokines than BEAS-2B cells. IGF2BP3 knock down increased *CCL5* and *CCL24* mRNA levels at baseline (*P* = 0.0004 and *P* = 0.039, respectively), and those of *CCL26* upon IL-4 stimulation (*P* = 0.0459). IL-13-driven responses were more pronounced, with increases in *CCL24* (*P* = 0.0099) and *CCL26* (*P* = 0.0462) upon IGF2BP3 reduction.

**Figure 4.**
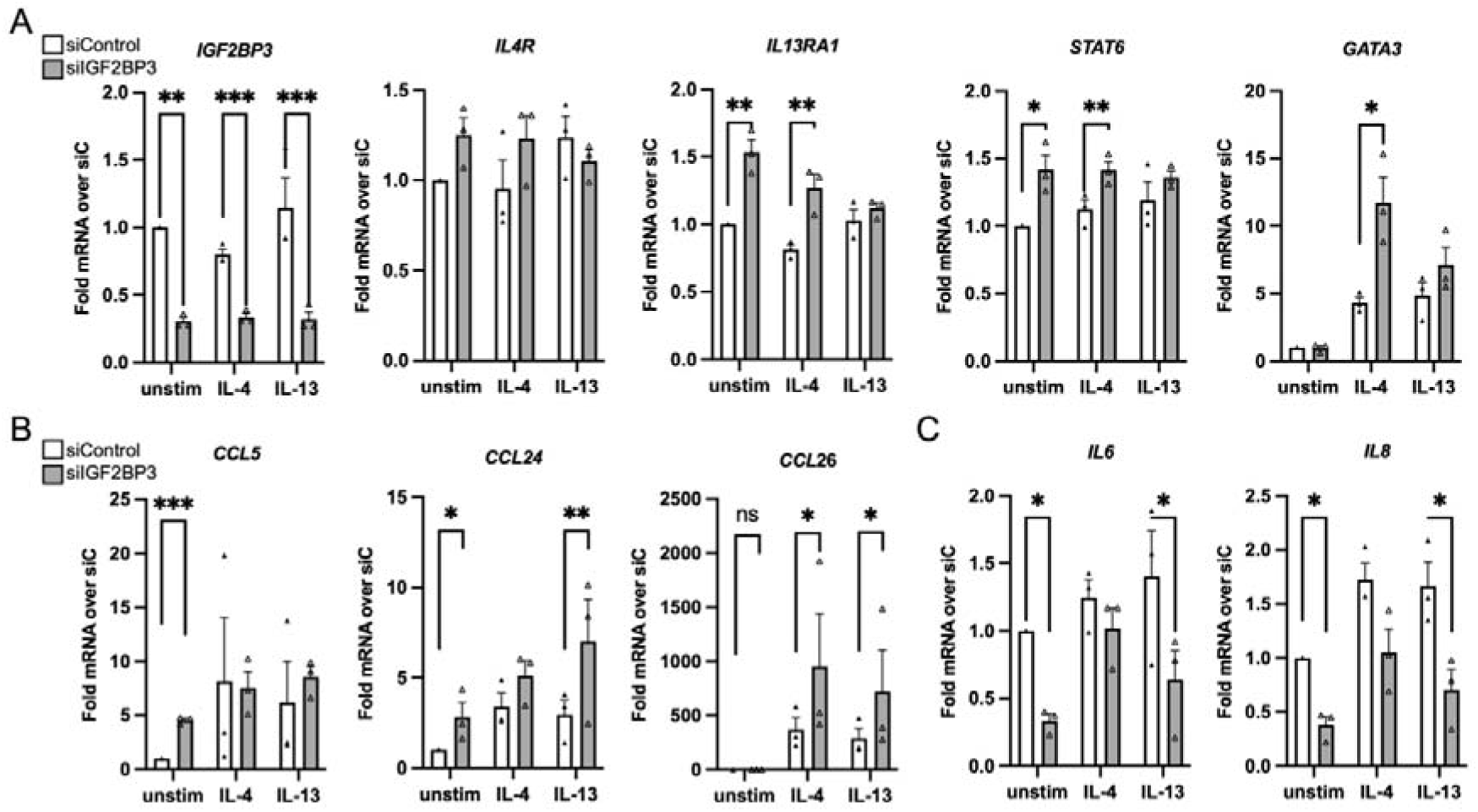
Decreasing IGF2BP3 levels increases IL-13/IL-4-driven transcriptional responses. RT-qPCR results of primary BECs transfected with siRNAs against IGF2BP3 (siIGF2BP3) or scrambled control (siControl) and stimulated with IL-13 or IL-4. A: RT-qPCR results for IGF2BP3 knock down and the major signalling mediators of IL-13 and IL-4. B: RT-qPCR results for chemokines. C: RT-qPCR results for pro-inflammatory *IL6* and *IL8* mRNAs. Data (n=3) depicted as mean ±SEM. One-tailed ratio t-tests. * *P* ≤ 0.05; ** *P* ≤ 0.01. unstim: unstimulated (referring to vehicle stimulated).

We also determined the effects on the mRNA levels of classically related ‘type 1’ mediators *IL6* and *IL8*. Figure 4C shows that IGF2BP3 knock down alone decreased both *IL6* and *IL8* mRNA expression (*P* = 0.0114 and *P* = 0.0267, respectively). While IL-4 stimulation appeared to even these differences, IL-13 stimulation further downregulated *IL6* (*P* = 0.0206) and *IL8* (P = 0.0256) levels in an IGF2BP3-dependent manner. Thus, silencing of IGF2BP3 in primary human bronchial epithelial cells alters their IL-13/IL-4 responses, skewing them towards a ‘type 2-promoting’ phenotype.

### IGF2BP3 depletion augments IL-13-driven effects on steady mRNA levels genome-wide

Our data pointed to a stronger and more consistent effect of IGF2BP3 depletion on the increase of IL-13-driven responses in BEAS-2B cells (Figure 3) and primary cells (Figure 4). IGF2BP3 modulation can lead to changes in steady, i.e. total, mRNA levels as well as their translation (21, 34, 35). Thus, we investigated the effect of IGF2BP3 depletion at the transcriptome level in different subcellular compartments using subcellular fractionation and RNA-sequencing (Frac-seq) (24, 36) using our constitutively depleted BEAS-2B cell lines stimulated with vehicle or IL-13 for 24 hours (Figure 5A).

**Figure 5.**
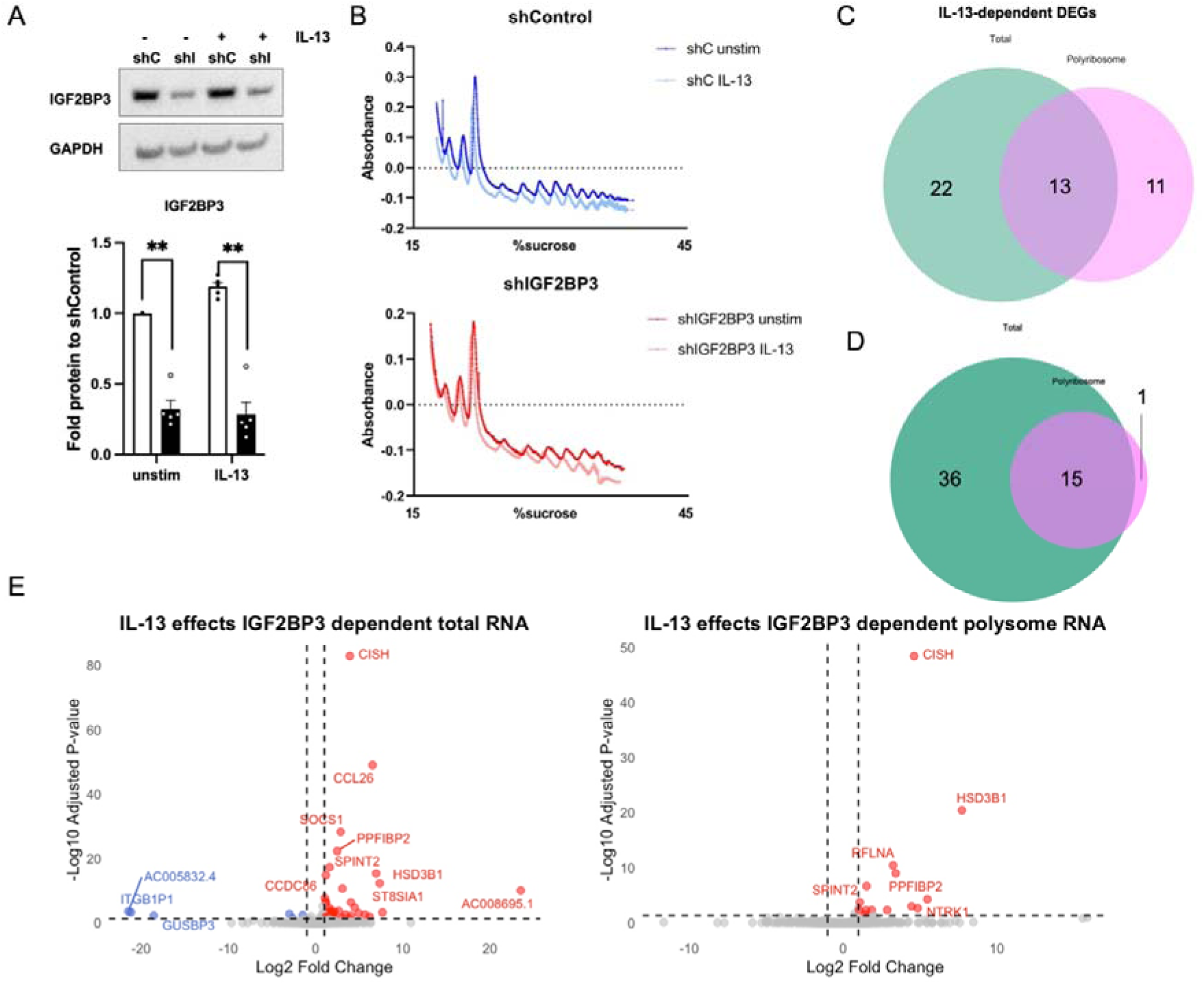
IGF2BP3 depletion modifies the effects on IL-13 genome-wide. A: representative western blot of our BEAS-2B constitutively depleted cell lines and quantification underneath (n=5). Paired two-tailed ratio t-test, data represented as mean ±SEM. B: Representative polyribosome profiles of BEAS-2B cell lines stimulated or not with IL-13. C: Venn diagram showing the similarities and differences of IL-13 stimulation in differentially expressed genes in Total and differentially bound genes in Polyribosome. D: Venn diagram showing the similarities and differences upon integrating the effects of IGF2BP3 depletion on IL-13 stimulation in differentially expressed genes in Total and differentially bound genes in Polyribosome. E: Volcano plots showing that IGF2BP3 depletion led to augmented effects of IL-13 stimulation, mainly in steady mRNA levels (Total, left plot). unstim: unstimulated (referring to vehicle stimulated).

We sequenced different subcellular factions, with ‘Total’ representing steady mRNA levels and ‘Polyribosome’ mRNAs associated with 5 ribosomes or more, i.e. mRNAs bound to the translational machinery and more likely to undergo translation (Figure 4B). IL-13 stimulation modified the mRNA levels of 35 mRNAs (p-adjusted < 0.05, Supplementary Table 2) and 24 mRNAs in the polyribosome fraction (p-adjusted < 0.05, Supplementary Table 3), of which 13 coincided with the changes observed in Total (Figure 4C). Thus, IL-13 exerts different effects in total vs polyribosome-bound mRNAs.

We also analysed the effects of decreasing IGF2BP3 levels (Supplementary Figure 3). IGF2BP3 preferentially modulated total mRNA levels (96 mRNAs, p-adjusted < 0.05, Supplementary Table 4) vs polyribosome binding of mRNA (20 mRNAs, p-adjusted < 0.05, Supplementary Table 5). For those mRNAs differentially regulated by IGF2BP3, we found increased 5’UTR and 3’UTR lengths as compared with the genome-wide expected distribution (Supplementary Figure 3).

Consistent with these separate findings, we modelled through DESeq2 analysis the effect of IGF2BP3 depletion on IL-13 stimulation (37). We found that the effects of IGF2BP3 on IL-13-dependent changes happened mostly at the steady mRNA level (Figure 5D, Supplementary Table 6). Increasing IGF2BP3 reverted some of these changes (Supplementary Figure 4). Of the 51 mRNAs that were differentially expressed, 15 were overlapping with polysome, which only had one discrete mRNA regulated (ENSG00000104332, corresponding to *SFRP1*, Supplementary Table 7). Thus, IGF2BP3 depletion increases the effects on mRNA expression triggered by IL-13 (Figure 5E).

### IGF2BP3 is upregulated in airway epithelium in type 2 disease characterised by an IGF2BP3-dependent IL-13 driven signature

Together, our data pointed towards a model where IGF2BP3 fine-tunes IL-13-driven responses. We then set out to contextualise the role of IGF2BP3 in airway epithelium from patients with type 2 disease, a prototype of IL-13-driven pathology. In addition to IGF2BP3 being upregulated in severe asthma *vs* healthy controls (24), we found it increased in atopic asthma vs healthy non-atopic controls (38). IGF2BP3 levels were also higher during virus-induced exacerbations as compared with post-exacerbation (39) (Figure 6A), virus-induced exacerbations in asthma characterised by a type 2 signature (40). Thus, IGF2BP3 appears increased in the airway epithelium of type 2 disease and type 2-driven responses.

**Figure 6.**
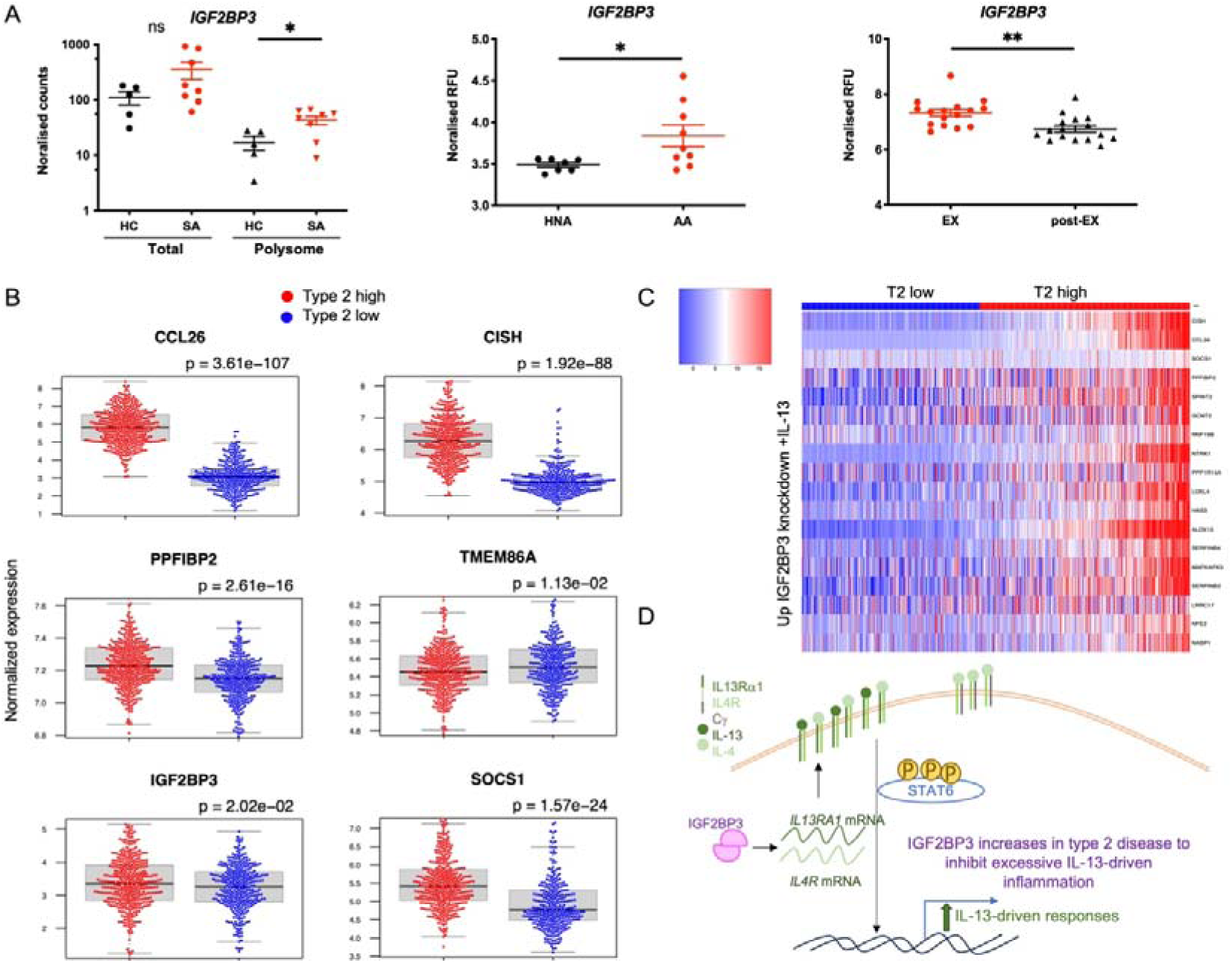
IGF2BP3 is increased in type 2 disease and associates with an IL-13-dependent signature in type 2 high asthma. A. Left: normalized expression levels of *IGF2BP3* mRNA in severe asthma (SA) vs healthy control (HC) primary BECs at total and polyribosome-bound levels; centre: *IGF2BP3* normalized microarray expression data comparing airway epithelial cells from healthy non-atopic (HNA) and atopic asthma (AA) children; right *IGF2BP3* normalized microarray expression data comparing nasal lavage samples from children during Picornavirus-induced exacerbation (EX) vs 7-14 days post-exacerbation (post-EX). B. Normalized count levels comparing type 2 high vs type 2 low nasal brushings for genes that showed IGF2BP3-dependent IL-13-modulation in our Frac-seq data (Figure 5E). C: Heatmap showing the normalized expression values of the common mRNA signature between differentially expressed genes in the GALA II cohort and our Frac-seq IGF2BP3 and IL-13-dependent DEGs. D. Model for IGF2BP3 effects on type 2 responses. Decreasing IGF2BP3 levels leads to increased stability of *IL13RA1* and *IL4R* mRNA and increased surface expression of IL4R and IL13Rα1, leading to increased STAT6 phosphorylation and increased IL-13 (and possibly IL-4) transcriptional responses, contributing to an overall increased type 2 response. In disease, IGF2BP3 is upregulated potentially to supress excessive T2 responses.

To test the relevance of IGF2BP3/IL-13 mediated changes in disease, we analyzed 695 nasal airway epithelial cell brushings from children (254 healthy controls and 441 asthmatics) from the GALA II cohort (41). We found several candidates of the IGF2BP3- and IL-13-dependent signature from Figure 5E differentially expressed between type 2 low and type 2 high patients (Figure 6B), and *IGF2BP3* also increased in type 2 high vs type 2 low patients, similarly to the suppressor of cytokine signalling *SOCS1*.

We next investigated the pattern of expression of IL-13-induced IGF2BP3-dependent mRNAs from Figure 5E in this cohort. Analysing the differentially expressed genes in common between our IL-13-induced IGF2BP3-dependent Frac-seq total mRNA levels and the expression levels of differentially expressed genes in GALA II, we found 20 candidates. Their expression pattern showed an enrichment for genes upregulated upon IGF2BP3 depletion and IL-13 stimulation to be increased in type 2 high vs type 2 low asthmatics (Figure 6C), serving as a potential IGF2BP3-dependent IL-13-induced signature in type 2 disease.

Considering our findings, we propose a model where reduction of IGF2BP3 increases the surface expression of Type II receptors, concomitant signalling and transcriptional responses of mainly IL-13-, and possibly IL-4-driven, mediators in human airway epithelium (Figure 6D). IGF2BP3 levels are increased in airway epithelium in type 2 disease, where it may be part of a feedback mechanism to reduce IL-13-driven inflammation.

## Discussion

Here we demonstrate that IGF2BP3 is expressed in human adult airway cells, where it inhibits IL-13/IL-4 signalling and is upregulated in type 2 disease. We initially found IGF2BP3 enriched in human airway epithelium from healthy donors. In human bronchial epithelial cells, IGF2BP3 co-immunoprecipitated *IL4R* and *IL13RA1* mRNAs, which are methylated. Decreasing IGF2BP3 levels led to increased mRNA half-live and surface expression of IL-4R and IL-13Rα1, as well as IL-4- and IL-13-dependent STAT6 phosphorylation. In turn, this led to increases in IL-13- and IL-4-driven transcriptional responses, particularly IL-13-dependent alarmin and cytokine secretion. We confirmed this type 2-skewing in primary human bronchial epithelial cells and further demonstrated that IGF2BP3 depletion increases the effects of IL-13-driven steady mRNA changes genome wide. IGF2BP3 is increased in airway epithelium from multiple cohorts of type 2 disease, where an IL-13-driven and IGF2BP3-dependent gene signature is overrepresented in type 2 high vs type 2 low asthma. Together, our findings suggest that IGF2BP3 is a new biomarker of type 2 disease, where its upregulation may reflect an attempt to dampen IL-13-driven pathology.

Since it was initially described, research on the effects of IGF2BP3 have mainly focused on its role as an oncogenic factor, as the established paradigm was that IGF2BP3 is mainly expressed in foetal tissue (16). Our findings are compatible with the previous roles described for IGF2BP3 as an oncogenic factor. Beyond the classical view of type 2 immunity as a suppressor of ‘anti-tumoral’ type 1 responses (42), type 2 immunity is led by epithelial alarmins which have been demonstrated as anti-tumorigenic (43, 44). In our BEAS-2B cell model, IGF2BP3 depletion led to increases in IL-13-dependent TSLP secretion (Figure 3C), which can block carcinogenesis early in breast cancer (45). Thus, further understanding of IGF2BP3 as a type 2 immune modulator in oncogenesis may open new therapeutic opportunities, particularly since the recent development of a potent small molecule inhibitor against IGF2BP3 (46).

Mechanistically, IGF2BP3 can exert myriad functions, including the regulation of mRNA stability, storage, or translation (18, 25, 30). Our data showed that IGF2BP3 depletion caused a mild increase in IL-4R and IL-13Rα1 surface expression (Figure 2D). These changes seemed to accompany an increase in their encoding mRNA half-lives, suggesting that IGF2BP3 regulates these mRNAs, at least in part, at the stability level (Figure 2D). Considering that IGF2BP3 is an m6A reader (21) and that these mRNAs were found to co-precipitate with an anti-m6A-antibody (Figure 2E), we also hypothesise that part of the molecular effects of IGF2BP3 rely on binding *IL4R* and *IL13RA1* methylated mRNAs and decreasing their stability. It is also possible that IGF2BP3 is modulating localized translation of *IL4R* and *IL13RA1* transcripts (25), an effect that will not be observable using polyribosome profiling. Intriguingly, we found IGF2BP3 expression more localized on the apical side of lung epithelium (Figure 1B) and it is thus possible that it may exert a subcellular location-dependent function. We did not find increased mRNA levels or binding to polyribosomes of *IL4R* and *IL13RA1* upon IGF2BP3 depletion in our Frac-seq data (Supplementary Tables 3 and 4), but we did however observe increases in *IL13RA1* mRNA in primary bronchial epithelial cells upon IGF2BP3 depletion (Figure 4A) where *IL13RA1* mRNA stability may be altered more profoundly than on BEAS-2B cells. Our data show the complementarity of our cell models, where effects in primary bronchial epithelial cells may better relate to IGF2BP3 role *in vivo,* and BEAS-2B cells serve as a model to work out the molecular mechanism.

Indeed, we observed differences between our BEAS-2B cells vs primary cells, including the chemokines they express (e.g. only *CCL26* was detectable in both). For example, *IL6* and *IL8* appeared downregulated in primary bronchial epithelial cells (Figure 4C), consequent with a change in their profile towards a ‘type 2’ and ‘anti-type 1’ skewing. In IGF2BP3-depleted BEAS-2B cells secreted more IL-8 in the presence of IL-13 (Figure 2C). We take these observations as a limitation of our experimental conditions, and they highlight the need for using the right cellular models – but they do not detract from our findings showing that depleting IGF2BP3 causes an increase in IL-13-driven effects.

Despite the above limitations, our bronchial epithelial cell model enabled us to find a group of 20 mRNAs that changed upon IL-13 stimulation in an IGF2BP3-dependent manner and whose expression appeared overrepresented in nasal samples from type 2 high vs type 2 low patients (Figure 6C). Intriguingly, we found that IGF2BP3 is increased in the airway epithelium of multiple cohorts with type 2 driven disease (Figure 6A, 6B). These results mirror our previous work where we demonstrated that miR-31 and miR-155 inhibited IL-13 signalling, but were increased in ulcerative colitis, an archetypal type 2 colonic disease (13). Back then, we proposed that these microRNAs may be part of the inhibitory loop to dampen type 2 inflammation. Here, we hypothesise that the increase in IGF2BP3 levels in type 2 disease may be part of the feedback mechanism to halt IL-13-driven pathology. Indeed, further increasing IGF2BP3 led to depletion of IL-13- (and IL-4)-dependent expression of *CCL26* and *SOCS1* mRNAs (Supplementary Figure 4), top candidates in our Frac-seq data. These type of feedback mechanisms are well established at the transcriptional level, such as increases in the levels of suppressor of cytokine signalling (SOCS) transcription factors upon cytokine stimulation (47), asthma (48) or ulcerative colitis (49). MicroRNAs regulate IL-13 responses post-transcriptionally (50) but the role of RBPs remains more obscure in this pathway. RBPs are known inhibitors of pro-inflammatory responses (14) and type 2 responses, although less investigated, may be no exception.

We are also aware that the increases in *IL4R* and *IL13RA1* mRNA levels and their encoded proteins are subtle fold changes, concomitant with a mild increased IL-13/IL-4-dependent STAT6 phosphorylation (Figure 3A). These subtle effects could translate into increased transcriptional responses, considering their ‘cascading effect’ in binding to promoter sequences and leading to more pronounced effects on IL-13-and IL-4-driven changes mRNA expression (Figures 3B, 4A and 4B). These changes were further confirmed at the genome-wide level, where we found IL-13-driven changes in steady mRNA expression to be further increased or decreased in IGF2BP3 depleted cells (Figure 5E). An alternative hypothesis is that IGF2BP3 may modulate other key factors within this pathway, for example *GATA3* (Figure 3B and 4A). GATA3 is widely known as an essentially factor in promoting the differentiation of Th2 cells (32); although it has been previously observed in bronchial epithelium (51), GATA3 function in airway epithelial cells remains poorly understood. Further exploration of balances between GATA3, STAT6 and IGF2BP3 levels and effects will shed more light into this possible type 2 circuit in lung epithelium. Considering the increased IGF2BP3 levels we observed in type 2 disease and our IL-13-driven and IGF2BP3-dependent signature, we propose IGF2BP3 as a new modulator of IL-4-and IL-13-driven homeostasis that is broken in type 2 disease.

In conclusion, we show for the first time that IGF2BP3 has immunomodulatory roles in human healthy airway cells and that is upregulated in type 2 disease, where it may attempt to counter-balance IL-13-driven pathology. To our knowledge, we show for the first time that RNA binding proteins can modify type 2 responses and are dysregulated in type 2 disease, positioning RNA biology at the core of IL-13/IL-4-driven responses and pathology.

## Methods

### Cell culture and cell lines

BEAS-2B immortalised bronchial epithelial cells were cultured in RPMI 1640 medium (ThermoFisher Scientific) supplemented with 10% foetal bovine serum (Merck Sigma-Aldrich). Primary bronchial epithelial cells (BECs) were commercially sourced (PromoCell), grown and expanded in growth factor-supplemented medium (PromoCell) on collagen-coated plates. Prior to stimulation, BEAS-2B cells were serum-starved overnight in RPMI with 2% FBS. All cells were maintained at 37°C in a 5% CO_2_ atmosphere and were passaged upon reaching 70-80% confluence. All datasets containing ‘unstimulated cells’ refer to vehicle-treated cells, we have maintained ‘unstimulated’ as nomenclature for ease of reading.

shRNAs (shControl or shIGF2BP3 from Origene) were sub-cloned between MluI and EcoRI sites of pLVTHM (Addgene, kindly donated by Prof Didier Trono). HEK293 T cells (kindly gifted by Prof Neil, King’s College London) at 90% confluence were transfected using Superfect (Qiagen) with lentiviral packaging vectors (psPAX2 and pMD2.G) and the pLVTHM vector. Virus-containing supernatants were collected 48 and 72 hours post-transfection, centrifuged to remove debris, and used to transduce BEAS-2B cells. BEAS-2B cells were plated at 50% confluence, cultured with 8μg/ml polybrene for 30 minutes and infected with lentivirus. Transduction was repeated after 24 hours. Post-transduction, cells were flow sorted for GFP+ cells in a BD Aria FACS cell sorter at the BRC Flow Core (GSTT).

### Cell transfection

siRNAs (Silencer Select, Darmacon) were used at a final concentration of 25nM. Transfection was carried out using Interferin (Polyplus) in accordance with the manufacturer’s protocol. IGF2BP3 overexpression was accomplished transfecting BEAS-2B cells with Lipofectamine 2000. Cells were incubated in the transfection mixture for 4 hours, after which the medium was replaced with BEC medium. Cells were stimulated with 100ng/ml IL-13 24 hours later.

### Public datasets

Figure 1A was taken from the Human Protein Atlas, www.proteinatlas.org, (28). Image available at the following URL: V24.proteinatlas.org/ENSG00000136231-IGF2BP3/tissue/bronchus#img. Figure 1C was taken from the GTEx Portal on 05/28/25.

Graphs corresponding to IGF2BP3 effects in Total and Polyribosome (Fig EV3) were taken from ShinyGO (https://bioinformatics.sdstate.edu/go/) (52). Figure 6 data and code corresponds to GSE85214 (24), GDS3711 (38), GDS4424 (39) and GSE152004 (41).

### Flow cytometry

shControl and shIGF2BP3 cells were washed twice with PBS and centrifuged at 300g for 5 minutes at 4°C. Cells were then resuspended in 200 μl PBS, and fluorescently labelled antibodies were added and incubated in the dark at room temperature for 20 minutes (IL-4Rα, dilution 1:200 of clone G077F6; IL-13Rα1, dilution 1:200 of clone SS12B, both antibodies PE labelled and from Biolegend). Upon washing, cells were acquired on a FACS Canto (BD Biosciences) and analysed using FlowJo software (TreeStar). For flow cytometry analysis, unstained samples and flurescence minus one (FMO) control were employed to set gates.

### Reverse transcription and quantitative PCR (RT-qPCR)

RNA extraction was carried out using TRIzol (ThermoFisher Scientific) or TRIzol LS (Merck Sigma-Aldrich). The extracted RNA was then reverse transcribed using RNAse H Minus Reverse Transcriptase, Ribolock, and random hexamers (ThermoFisher Scientific). Quantitative PCR (qPCR) was conducted with NEB Luna buffer (New England Biolabs) and TaqMan assays (Supplementary Table 1, ThermoFisher Scientific). Gene expression levels were analyzed using the 2-^ΔCt^ method (53) and normalized to GAPDH (Primer Design).

### RNA stability assay

BEAS-2B constitutive cell lines (shControl and shIGF2BP3) were grown in 6-well plates overnight. The following day, they were treated with 5µg/ml actinomycin D to block gene transcription for different time intervals. RNA was subsequently extracted quantified using Nanodrop and assessed by RT-qPCR as above. Gene expression levels were normalized to GAPDH (Primer Design).

### Western blot

Cells were lysed on ice using polysome lysis buffer (0.5% NP40, 20 mM Tris HCl pH 7.5, 100 mM KCl, 10 mM MgCl2) supplemented with fresh protease inhibitor cocktail (Cell Signaling). The lysate was separated on SDS-polyacrylamide gels and subsequently transferred to a nitrocellulose membrane (BioRad Laboratories). The membrane was blocked in blocking buffer (Amersham Biosciences) for 60 minutes at room temperature, then incubated overnight at 4°C with primary antibodies (IGF2BP3: Santa Cruz sc-390639, GAPDH: Santa Cruz sc32233, STAT6: CST 5397S, Phospho-STAT6: CST 9361S). Afterwards, the membrane was incubated for 60 minutes at room temperature with the appropriate horseradish peroxidase-conjugated anti-rabbit or anti-mouse IgG secondary antibodies (Dako). All washes and antibody dilutions were performed in TBS-0.05% Tween. Immunoreactive proteins were detected using the ECL immunodetection system (ThermoFisher Scientific) and visualized with the ChemiDoc MP Imaging System (BioRad), with analysis performed using ImageLab (BioRad) software.

### Cytokine and chemokine detection

Our shControl and shIGF2BP3 constitutive cell lines cells were seeded and 24 h later starved overnight, followed by stimulation with 100 ng/ml IL-4, 100 ng/ml IL-13 or vehicle. 24 hours later the supernatants were collected for cytokine analysis using a U-PLEX biomarker assay kit (Meso Scale Discovery) as per manufacturer’s instructions. Briefly, a set of 7 standards was created through 4-fold serial dilutions. Biotinylated antibody was mixed with the assigned linker and incubated at room temperature for 30 minutes, followed by addition of a stop solution at room temperature for 30 minutes. 600 μl of each linker-coupled antibody solution were combined into a single tube, and the solution was brought up to a 1X concentration by increasing its volume to 6 ml. 50 μl of the 1X multiplex coating solution were added to each well; the plate was sealed and shaken for 1 hour at room temperature and washed three times with wash buffer. 25μl of assay diluent and 25 μl of the prepared standards or samples were added and incubated at room temperature for 1 hour with shaking. The plate was washed three more times with wash buffer, and 50 μl of the detection antibody solution was added to each well. The plate was sealed and incubated at room temperature with shaking for 1 hour. The wash steps were repeated three more times before the results were read using a 1300 MESO QuickPlex, following the manufacturer’s instructions.

### RNA immunoprecipitation (RIP)

RNA immunoprecipitation (RIP) was performed using 2.1μg of IGF2BP3 and corresponding IgG antibodies (Merck Sigma-Aldrich: 03-198) and 5μg Anti-N6-methyladenosine (m6A) antibody (ab151230) and IgG_1_ (ThermoFisher Scientific) per 1 million cells. BEAS-2B cells were lysed on ice in RIP lysis buffer (100mM KCl, 25 mM Tris pH 7.4, 5 mM EDTA, 0.5 mM DTT, 0.5% NP40) supplemented with 100 U/mL Ribolock RNase inhibitor (ThermoFisher Scientific) and protease inhibitor cocktail (Cell Signaling Technology). Ten percent of the lysate volume was retained as input. For antibody binding, Dynabeads Protein A magnetic beads (ThermoFisher Scientific) were used, and the antibody-bead complex was incubated on a rotating shaker for 10 minutes at room temperature. The cell lysates were then immunoprecipitated with the antibody-bead complex on a rotating shaker for 10 minutes. RNA was extracted from the beads using TRIzol (ThermoFisher Scientific), and RNA presence was measured by RT-qPCR.

### Polyribosome profiling

Polyribosome profiling was performed as previously described (24). Briefly, constitutive cell lines were stimulated with 100 ng/mL IL-13, and 500 μg/mL cycloheximide (Merck Sigma-Aldrich) were added 10 minutes prior to lysis. Supernatants were carefully collected and reserved for cytokine analysis. Cells were lysed on ice for 7 minutes in polyribosome lysis buffer (0.5% NP40, 20 mM Tris HCl pH 7.5, 100 mM KCl, 10 mM MgCl_2_, 1X protease inhibitor cocktail (Cell Signaling, 5871S), 100 μg/mL cycloheximide) and then passed through a 23G needle three times. Cytoplasmic extracts were centrifuged at 8000 rpm for 7 minutes at 4°C. Ten percent of the cell lysate was reserved for cytoplasmic (Total) RNA extraction, and the remaining lysate was carefully loaded onto 15-45% sucrose gradients and ultracentrifuged using a Hitachi ultracentrifuge. The preparation of gradients, the reading of polyribosome profiles, and the extraction of individual polyribosome peaks were performed using a Gradient Station (BioComp Instruments, Fredericton, NB, Canada) equipped with a Triax. Polyribosome profiles were generated by measuring the absorbance at 260 nm of the spun gradients immediately after ultracentrifugation. Fractions corresponding to each ribosomal peak were isolated and extracted using TRIzol LS (Sigma-Aldrich) according to the manufacturer’s instructions. RNA from fractions P5 (6 ribosomes) onwards was pooled to form the polyribosome fraction, which was sequenced along with the total RNA fraction.

### Library preparation and RNA-sequencing

Libraries were prepared using the NEBNext Ultra II Directional RNA Library Prep Kit for Illumina (New England Biolabs) with poly(A) mRNA selection and dual-index primers. An input of 10 ng RNA was used. Sequencing was performed on a NovaSeq6000 (Illumina) with 150bp paired-end reads to a depth of 40 million reads.

### RNA sequencing

Data analysis was conducted using the Cancer Genomics Cloud (CGC) (54). Quality control of the sequencing data was performed with FastQC (55), and adapter sequences were removed using Trimmomatic (56). Alignment to the genome (v43) was carried out with Salmon, utilizing the Salmon index and Salmon quant (57).

Differential gene expression analysis was conducted in R Studio using DESeq2 (37), along with all downstream data visualization. The data was generated from five biological replicates, with gene expression levels reported as log_2_FoldChange. A p-adjusted value (p-adj) of less than 0.05 was considered significantly regulated.

### Statistical analysis

GraphPad Prism and R Studio were used for data analysis and graph generation. Data are presented as mean ± standard error of the mean (SEM). Normality was assessed using the Shapiro-Wilk test. Statistical significance was indicated by p-values: p < 0.05 (\**), p < 0.01 (**), and p < 0.001 (\*\*\**). Statistical analyses of control vs stimulated cells, as well as shControl vs shIGF2BP3 were performed paired and two-tailed; data were either log transformed to perform t-tests or tested as ratio tests (which average the logarithm of the ratio condition/control and better test for differences between fold changes). Actinomycin D experiments were analysed using a one phase exponential decay as per (31).

## Acknowledgements

We are incredibly grateful to the authors of (41) for providing such detailed code of their work. This work was supported by King’s College London and King’s Health Partners Challenge Fund.

## Author contributions

P.K. and O.Z. performed the experiments and revised the manuscript; J.R. contributed with data analysis; E.P. contributed with experiments, analysis and manuscript writing; R.T.M-N. conceived and supervised the study, performed analysis and wrote the manuscript.

## Declaration of interests

R.T.M-N declares funding from GSK and AZ for studies not related with this current work.

## Data Availability Section

Datasets have been deposited in Gene Expression Omnibus GSE300481. A reviewer link is available upon request.

Our code will be available in GitHub https://github.com/rociotmartinez/IGF2BP3

## Notes

https://V24.proteinatlas.org/ENSG00000136231-IGF2BP3/tissue/bronchus#img

https://www.gtexportal.org/home/

https://www.ncbi.nlm.nih.gov/geo/query/acc.cgi?acc=GSE18965

https://www.ncbi.nlm.nih.gov/geo/query/acc.cgi?acc=GSE85216

https://www.ncbi.nlm.nih.gov/geo/query/acc.cgi?acc=GSM751969

https://www.ncbi.nlm.nih.gov/geo/query/acc.cgi?acc=GSE152004

### Summary of Updates

We have made the text neater and added supplementary materials to support our findings that IGF2BP3 inhibits IL-4/IL-13 signalling

