## Supplementary for "IGF2BP3 inhibits IL-13 and IL-4 effects in human airway epithelium and is dysregulated in type 2 disease"

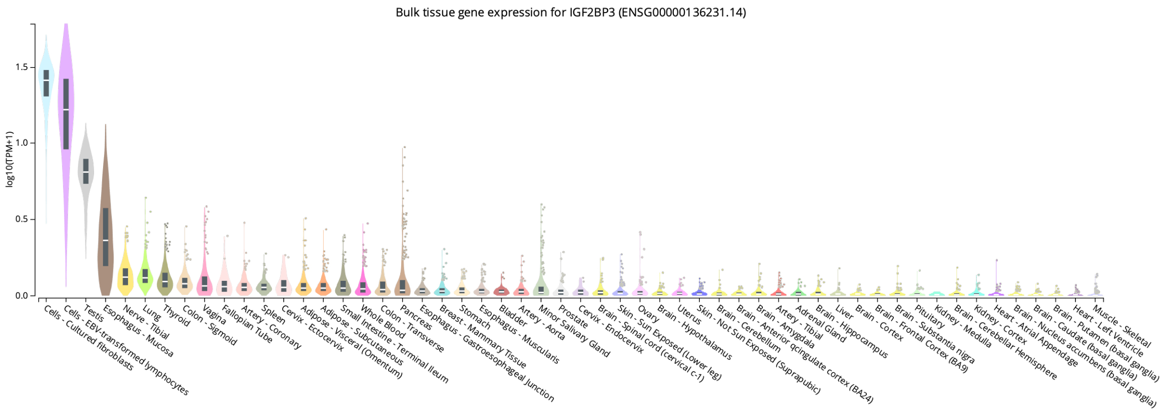


**Supplementary Figure 1. GTEx plot showing IGF2BP3 expression as per RNA-seq in their tissues analyzed.**
Data from https://www.gtexportal.org/home/ (Consortium, 2013)

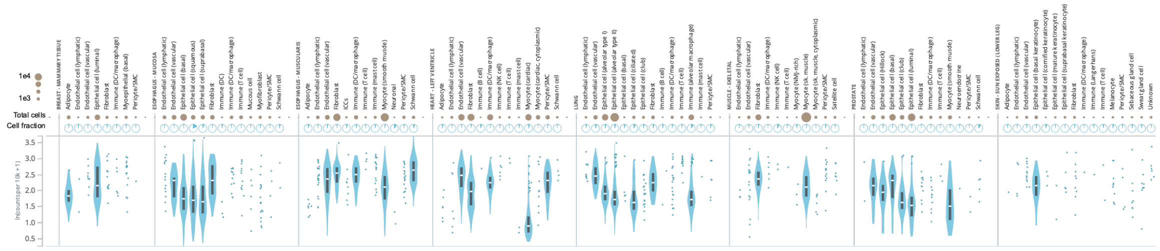


**Supplementary Figure 2. GTEx single cell expression showing IGF2BP3 expression per cell type and tissue, excluding zero counts.**
Data from https://www.gtexportal.org/home/ (Consortium, 2013)

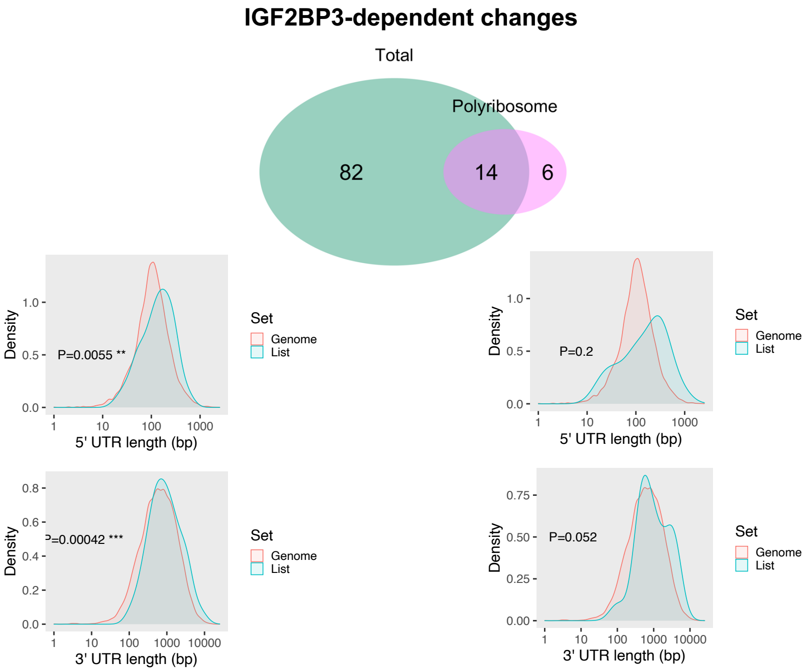


**Supplementary Figure 3. Effects of IGF2BP3 knock down on BEAS-2B cells**. IGF2BP3 depletion caused more effects on steady mRNA levels vs polyribosome binding (Venn diagram), where mRNAs appeared to have increased 5'UTR and 3'UTR lengths as compared with the genome-wide distribution. Differentially bound mRNAs upon IGF2BP3 did now show any statistically significant differences in their lengths as compared with the expected distributions genome-wide. Figures from https://bioinformatics.sdstate.edu/go/ (Ge *et al*, 2019)

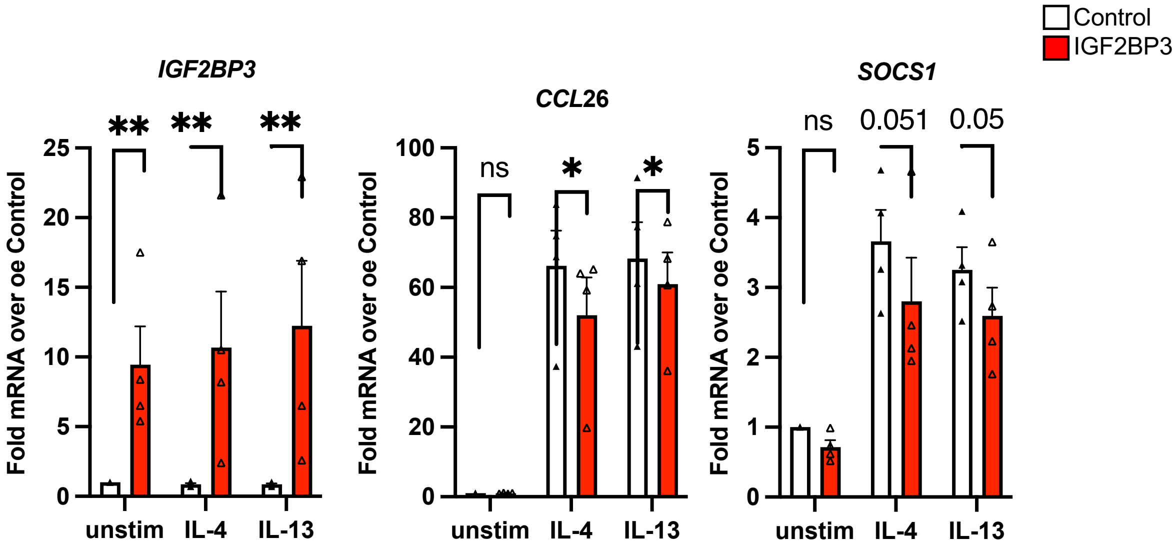


**Supplementary Figure 4. Overexpression of IGF2BP3 decreases IL-4- and IL-13-dependent mRNA expression.** BEAS-2B cells were transfected with a plasmid overexpressing IGF2BP3 or control, and exposed to vehicle (unstim), IL-4 or IL-13 for 24h. RNA was harvested and analysed by RT-qPCR.
